# A worldwide ENIGMA study on epilepsy-related gray and white matter compromise across the adult lifespan

**DOI:** 10.1101/2024.03.02.583073

**Authors:** Judy Chen, Alexander Ngo, Raúl Rodríguez-Cruces, Jessica Royer, Maria Eugenia Caligiuri, Antonio Gambardella, Luis Concha, Simon S. Keller, Fernando Cendes, Clarissa L. Yasuda, Marina K. M. Alvim, Leonardo Bonilha, Ezequiel Gleichgerrcht, Niels K. Focke, Barbara Kreilkamp, Martin Domin, Felix von Podewils, Soenke Langner, Christian Rummel, Roland Wiest, Pascal Martin, Raviteja Kotikalapudi, Benjamin Bender, Terence J. O’Brien, Benjamin Sinclair, Lucy Vivash, Patrick Kwan, Patricia M. Desmond, Elaine Lui, Gian Marco Duma, Paolo Bonanni, Alice Ballerini, Anna Elisabetta Vaudano, Stefano Meletti, Manuela Tondelli, Saud Alhusaini, Colin P. Doherty, Gianpiero L. Cavalleri, Norman Delanty, Reetta Kälviäinen, Graeme D. Jackson, Magdalena Kowalczyk, Mario Mascalchi, Mira Semmelroch, Rhys H. Thomas, Hamid Soltanian-Zadeh, Esmaeil Davoodi-Bojd, Junsong Zhang, Matteo Lenge, Renzo Guerrini, Emanuele Bartolini, Khalid Hamandi, Sonya Foley, Theodor Rüber, Tobias Bauer, Bernd Weber, Benoit Caldairou, Chantal Depondt, Julie Absil, Sarah J. A. Carr, Eugenio Abela, Mark P. Richardson, Orrin Devinsky, Heath Pardoe, Mariasavina Severino, Pasquale Striano, Domenico Tortora, Erik Kaestner, Sean N. Hatton, Donatello Arienzo, Sjoerd B. Vos, Mina Ryten, Peter N. Taylor, John S. Duncan, Christopher D. Whelan, Marian Galovic, Gavin P. Winston, Sophia I. Thomopoulos, Paul M. Thompson, Sanjay M. Sisodiya, Angelo Labate, Carrie R. McDonald, Lorenzo Caciagli, Neda Bernasconi, Andrea Bernasconi, Sara Larivière, Dewi Schrader, Boris C. Bernhardt

## Abstract

**Objectives:** Temporal lobe epilepsy (TLE) is commonly associated with mesiotemporal pathology and widespread alterations of grey and white matter structures. Evidence supports a progressive condition although the temporal evolution of TLE is poorly defined. This ENIGMA-Epilepsy study utilized multimodal magnetic resonance imaging (MRI) data to investigate structural alterations in TLE patients across the adult lifespan. We charted both grey and white matter changes and explored the covariance of age-related alterations in both compartments.

**Methods:** We studied 769 TLE patients and 885 healthy controls across an age range of 17-73 years, from multiple international sites. To assess potentially non-linear lifespan changes in TLE, we harmonized data and combined median split assessments with cross-sectional sliding window analyses of grey and white matter age-related changes. Covariance analyses examined the coupling of grey and white matter lifespan curves.

**Results:** In TLE, age was associated with a robust grey matter thickness/volume decline across a broad cortico-subcortical territory, extending beyond the mesiotemporal disease epicentre. White matter changes were also widespread across multiple tracts with peak effects in temporo-limbic fibers. While changes spanned the adult time window, changes accelerated in cortical thickness, subcortical volume, and fractional anisotropy (all decreased), and mean diffusivity (increased) after age 55 years. Covariance analyses revealed strong limbic associations between white matter tracts and subcortical structures with cortical regions.

**Conclusions:** This study highlights the profound impact of TLE on lifespan changes in grey and white matter structures, with an acceleration of aging-related processes in later decades of life. Our findings motivate future longitudinal studies across the lifespan and emphasize the importance of prompt diagnosis as well as intervention in patients.

## INTRODUCTION

Epilepsy is a common and debilitating neurological disease affecting over 50 million individuals worldwide, of which a substantial proportion suffers from functional deficits and markedly compromised quality of life.^1^ Temporal lobe epilepsy (TLE) is the most common pharmaco-resistant epilepsy in adults associated with mesiotemporal pathology.^2^ While traditionally considered a prototypical “focal” epilepsy, increasing evidence suggests that atrophy and white matter alterations are not limited to the mesiotemporal region, but affect widely distributed grey and white matter systems.^3,4^ Single and multi-site morphometric studies have reported significantly reduced grey matter volume in temporal, frontal, and centroparietal cortices as well as subcortical structures such as the thalamus and amygdala.^5–7^ Moreover, there is a compromise of white matter microstructure and architecture of numerous fiber tracts, with pronounced effect sizes in limbic and subcortico-cortical systems.^8^ Alterations have been shown to follow structural connectivity patterns between distributed regions and the mesiotemporal epicentre, and impact inter-regional morphometric covariances.^9,10^ As such, TLE is now increasingly recognized as a “network” disorder accompanied by profound changes in cortico-subcortical grey matter morphology and changes in related white matter compartments.^3,4,11,12^

There is evidence that TLE is not a static condition, but one that shows progression over time. Advanced age is a critical risk factor for cognitive decline, particularly in TLE where the majority of patients older than 55 years have present with memory and language deficits that may meet the criteria for mild cognitive impairment.^13^ In addition to cognitive decline, TLE patients exhibit accelerated bilateral cortical thinning beyond the mesiotemporal region.^14,15^ A recent meta-analysis of structural magnetic resonance imaging (MRI) studies suggested that TLE is associated with cumulative atrophy exceeding typical aging-related processes, but most constituent studies were limited by the low quality of evidence and heterogeneous approach to controlling for age-related effects.^16^ Even fewer studies have investigated progression of effects on white matter despite the role of this compartment in seizure propagation. Despite some evidence for age as a key determinant of global white matter volume changes, aging-related effects on white matter microstructure remain underexplored.^17^

Thus, whilst these prior investigations provide initial evidence for age-modulated changes in brain structure in TLE, findings are limited in several ways. First, most research has focused on grey matter morphology,^16^ not addressing white matter changes nor the covariance between grey and white matter alterations. Such studies would be needed to understand aging effects on brain integrity and to identify pathology affecting white and grey matter in epilepsy. Second, research into aging effects in TLE has been limited by focusing on only small cohorts and has been mainly restricted to single sites.^15,18,19^ Such an approach may be limited in terms of reproducibility and generalizability and affected by cohort- and centre-specific idiosyncrasies.^5^ Third, age is generally considered a confounder for the more prevailing analyses of disease duration or between-group difference effects, and has therefore not been a key focus in assessing its direct relation to MRI measures in TLE in prior work.^16^ Here, we address these gaps by leveraging a multi-site TLE cohort aggregated by the ENIGMA-Epilepsy consortium to map lifespan changes in multimodal MRI measures of grey matter morphology and white matter microstructure.^6,8^ In addition to assessing grey and white matter changes independently, we examined their covariance to quantify the inter-regional and cross-compartment coupling of aging-related processes.

## METHODS

### Participants

We studied 769 TLE patients (455 females, mean ± standard deviation [SD] age = 38.2 ± 11.1 years, 430/339 with left/right-sided seizure focus, age of onset = 16.5 ± 12.0 years, disease duration = 21.3 ± 13.5 years) and 885 age- and sex-matched healthy controls (508 females, mean ± SD age = 36.0 ± 11.7 years) spanning an age range of 17-73 years from the ENIGMA-Epilepsy cohort. Summary clinical and demographic characteristics are presented in **Table S1**. Patients that had undergone surgery with Engel score I (n=216) are reported as well. TLE was diagnosed according to standardized International League Against Epilepsy criteria across 18 recruitment sites.^20^

### Multimodal MRI data processing and harmonization

Acquisition and scanner descriptions of both the structural T1-weighted (T1w) MRI and diffusion-tensor imaging (DTI) for neocortical and subcortical regions obtained from all participants are described in previous ENIGMA-Epilepsy studies.^6,8^ Images were independently processed on site using a standardized workflow (T1w: https://enigma.ini.usc.edu/protocols/imaging-protocols/, DTI: https://enigma.ini.usc.edu/protocols/dti-protocols/, https://enigma.ini.usc.edu/ongoing/dti-working-group/)^5^, which allowed for the measurement of cortical thickness (CT) and subcortical volume (SV) based on T1w according to the Desikan-Killiany atlas, composed of 68 cortical and 16 subcortical regions.^21^ Diffusion MRI data were processed using TBSS to derive the tensor parameters fractional anisotropy (FA) and mean diffusivity (MD) for 38 unique white-matter tracts referenced from the JHU ICBM-DTI-81 atlas.^22^

### Statistical analysis

Analyses were performed in Matlab R2019b (The Mathworks, Natick, MA). To control for potential confounders such as scanner and site variability but to preserve biological covariates (age, sex, and diagnostic group), harmonization via ComBat was performed for DTI and T1w data.^23,24^ Patient measures were *z*-scored relative to controls and sorted into ipsilateral/contralateral to the seizure focus as per previous studies.^9,25^ We then used the openly accessible BrainStat toolbox for statistical inference and data visualization (http://brainstat.readthedocs.io).^26^

### a) Age-bisected cohort analysis

To preserve statistical power of this large dataset, while establishing age-dependent differences between TLE patients and controls, the cohort was first split via its median age across the entire sample (35 years), which also approximates to the median in TLE (37 years) and control (33 years) groups. This resulted in a “young” (< 35 years) and “older” (>= 35 years) cohort for both patients (n young/old=292/477) and healthy controls (n=481/405). Linear models highlighted CT, SV, FA, and MD effect size (Cohen’s *d*) profiles of young patients as compared to their age-equivalent healthy control cohort, and likewise of old patients and their age-equivalent control cohort. This illustrated the progression of differences between the two age groups. Group-by-age interactions analysis identified regions showing more marked age-dependent changes in TLE patients compared to controls, with participant ages treated categorically to identify only regions with the most robust age-dependent changes and to align with the prior median split analyses. All findings controlled for sex and were corrected for multiple comparisons using the false discovery rate (FDR) procedure.^27^

### b) Sliding age-window and covariance analysis

Age-sliding windows were utilized to stabilize trends and limit the influence of potential outliers.^28^ Though various window ranges can be constructed, a range of ±2 years from an age of interest (AOI) was chosen for presented analyses to best maintain granularity of trajectory perturbations while maximizing the number of patients. Sensitivity analyses were conducted to ensure preservation of trends across different age windows (**Fig. S3**). AOI and subsequent windows begin at 19 and end at 70 years of age; each window is slid incrementally by 1 year, resulting in 52 unique sub-cohorts. For each sub-cohort, structural/microstructural measures were calculated via normally distributed weights based on their age distance to the AOI. Finally, each sliding age-window was *z*-scored against the healthy control cohort to yield age-associated *z*-score trajectories across all cortical, subcortical, and white matter tract regions (**Fig. 1B**). Correlations of each region-associated age trajectory across the lifespan for CT, SV, FA, and MD were also calculated, with the top 10% and bottom 10% highlighted to identify region(s) for which their lifespan trajectories are most closely related.

**Figure 1.**
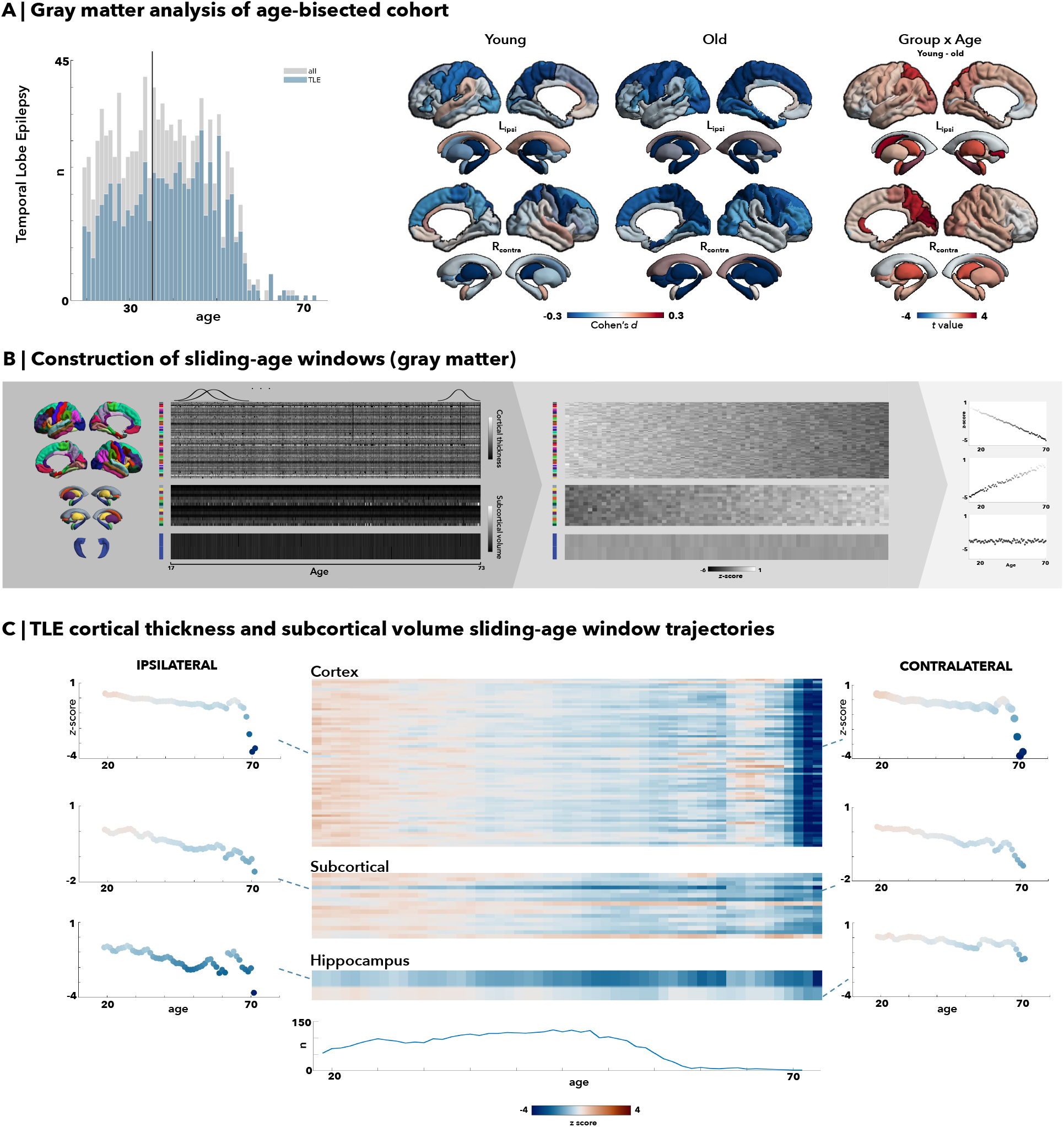
Grey matter morphometry via median-split analysis and sliding-age window trajectories in temporal lobe epilepsy (TLE). **(A)** Median age of 35 was chosen as the division into young and old cohorts. All right-sided temporal lobe epilepsy (TLE) patients were flipped, with results demonstrating significant left hippocampal reductions in both young and old age cohorts, and increasing occipital grey matter atrophy with age as seen in the old cohort. There are strong group by age interaction with significant subcortical regions and some cortical involvement in the parieto-occipital regions. **(B)** After z-scoring cortical thickness and subcortical volume measures, windows of ±2 years of the age of interest were chosen, and the mean values for cortical (68 total brain regions) and subcortical regions (12 subcortical grey matter regions and bilateral ventricles and hippocampi) following the Desikan-Killiany atlas were multiplied by normally distributed weights to yield a weighted average. Global weighted-average z-scores were then plotted across age to illustrate lifespan trajectory. **(C)** In temporal lobe epilepsy (TLE) patients, older age especially past age 55 seems to be correlated with an accelerated and pronounced decline in cortical thickness. The hippocampi and other subcortical regions also show a marked decreased volume with aging.

## RESULTS

### Grey matter morphometric lifespan analysis

Median split analysis showed an increased rate of ipsilateral reductions in CT and SV in TLE patients over adulthood as compared to changes seen in healthy individuals (**Fig. 1A**). Young TLE patients exhibited bilateral neocortical thinning in the precentral (*d*=-0.22), postcentral (*d*=-0.28), superior parietal (*d*=-0.27), and middle frontal (*d*=-0.27) regions compared to their age-matched healthy counterparts (p_FDR_<0.05). In the ipsilateral hemisphere, there was additional atrophy in the transverse temporal areas (*d*=-0.20, p_FDR_<0.05). The differences between patients and controls in the older TLE cohort showed almost identical overlap for atrophic regions relative to the results of the younger cohorts, but with further atrophy of the neocortex, particularly in the ipsilateral hemisphere. Additional areas of atrophy in the older age group included ipsilateral entorhinal (*d*=-0.26, p_FDR_<0.0005) and parahippocampal regions (*d* = -23, p_FDR_<0.005), as well as further bilateral inferior parietal (*d* ipsi/contra=-0.20/-0.18, p_FDR_<0.05) and contralateral occipital (*d*=-0.23, p_FDR_<0.005) regions. While the contralateral hemisphere presented with fewer significant regions, atrophy overall still included a similar pattern as seen in the younger TLE cohort with additional effects in bilateral frontal (superior frontal *d* ipsi/contral=-0.23/-0.21, caudal middle frontal (*d* ipsi/contra=-0.24/-0.24, p_FDR_<0.05) thinning. In total, 18 (*d* ipsi/contra=11/7) cortical regions of interest (ROI) significantly differed in the young TLE cohort, while 26 (*d* ipsi/contra=16/10) ROIs significantly differed in the older TLE cohort. Overall, the group-by-age interaction analysis showed that neocortical thinning in the ipsilateral superior parietal, contralateral precuneus, cuneus, and pericalcarine regions was significantly more marked in older patients than younger patients in comparison to their healthy control counterparts (p_FDR_<0.05).

There were significant ipsilateral hippocampal volume reductions in both young and older TLE cohorts compared to their healthy age counterparts (*d*=-0.60, p_FDR_<0.05e-15). Young TLE cohorts exhibited bilateral volume reductions in the thalamus (*d* ipsi/contra=-0.41/-0.29, p_FDR_<0.05) and pallidum (*d* ipsi/contra=-0.17/-0.19, p_FDR_<0.05). Older TLE cohorts showed more extensive, bilateral findings (p_FDR_<0.05), with greater volume reductions in the hippocampi (*d* ipsi/contra=-0.82/-0.67), thalamus (*d*=-0.29/-0.68), caudate (*d*=-0.41/-0.31), and ipsilateral amygdala (*d*=-0.26).

To discern trajectories of neocortical thinning over the lifespan (**Fig. 1B-C**), age-sliding windows were constructed, and cortical thickness was z-scored against controls for each region. Overall, there was a slow but progressive decline in mean cortical thickness from young adulthood to middle age, with mean z-scores decreasing to -1 by age 55 and dropping more steeply to -4 beyond age 60; this pattern was reflected across all cortical regions bilaterally. In contrast, subcortical volumes decreased linearly and less severely (ranging from *z*-scores of 0 to -2) as age advanced. This pattern remained consistent across all subcortical regions and the hippocampus. In particular, the volume of the ipsilateral hippocampus exhibited the most consistent decline, with substantially decreased *z*-scores compared to other cortical and subcortical regions. Furthermore, comparative sensitivity analyses of these age-dependent trajectories demonstrated much higher correlation with the TLE-mesial temporal sclerosis (MTS) sub-cohort (CT/SV *r*=0.88/0.91) compared to the TLE-non lesional (NL) sub-cohort (CT/SV *r*=0.68/0.70). There were little differences between the correlations of the left-sided TLE (CT/SV *r*=0.77/0.83) and right-sided TLE (CT/SV *r*=0.82/0.86) sub-cohorts.

### White matter microstructure lifespan analysis

Within the age-bisected cohort, young TLE patients exhibited smaller FA decreases compared to older TLE patients against their respective age-matched control cohort with a total of 19/21 significantly different ROIs (p_FDR_<0.05) within the younger TLE cohort, and 21/21 significantly different ROIs (p_FDR_<0.05) within the older TLE cohort. Most white matter tracts within the younger TLE cohort were significantly decreased, particularly in the fornix (FX: *d*=-0.37; fornix stria terminalis [FXST]: *d* ipsi/contra=-0.29/-0.33 p_FDR_<0.05), genu (GCC:-0.41, p_FDR_<0.05) and body of the corpus callosum (BCC: -0.43, p_FDR_<0.05), bilateral external capsule (EC: *d* ipsi/contra=-0.45/-0.53, p_FDR_<0.05), and cingulum (cingulate gyrus [CGC]: *d* ipsi/contra=-0.42/-0.47 ; cingulum adjacent to the hippocampus hippocampus [CGH]: *d* ipsi/contra=-0.3/-0.42, p_FDR_<0.05). All white matter tracts were significantly decreased in older TLE patients with even greater decline in FA values within the fornix (FX: *d* = -0.59, p_FDR_<0.05; FXST: *d* ipsi/contra=-0.45/-0.47, p_FDR_<0.05), genu (GCC: *d*=-0.61, p_FDR_<0.05) and body (BCC: *d*=-0.72, p_FDR_<0.05) of the corpus callosum, and cingulum (CGC: *d* ipsi/contra=-0.60/-0.61; CGH *d* ipsi/contra=-0.54/-0.57; p_FDR_<0.05). Overall, there was a clear decrease in FA values between older cohorts and younger cohorts, with significant age-effects across almost all white matter regions, suggesting a weakening of structural integrity of the fibre tracts and cumulative damage with age.

For MD, we observed a low to moderate increase in most tracts in young TLE cohorts when compared to their age-matched controls, totalling 14/21 bilaterally altered ROIs (p_FDR_<0.05). The fornix (*d*=0.41, p_FDR_<0.05), bilateral external capsule (EC: *d* ipsi/contra=0.31/0.32, p_FDR_<0.05), cingulum adjacent to the hippocampus (CGH: *d* ipsi/contra=0.33/0.30, p_FDR_<0.05), and the anterior and posterior corona radiata (ACR: *d* ipsi/contra=0.25/0.23; PCR: *d* ipsi/contra=0.27/0.26, p_FDR_<0.05) had the highest MD increases within the young TLE cohort. The ipsilateral corticospinal tract was the sole structure that had an unexpected decrease in MD (*d*=-0.65, p_FDR_<0.05). In the older TLE cohort, almost all (20/21 ROIs) white matter structures demonstrated larger increases as compared to the differences seen in the younger TLE cohort (p_FDR_<0.05), with the greatest increases in the corpus callosum (GCC: *d*=0.50; BCC: *d*=0.65, p_FDR_<0.05), corona radiata (ACR: *d* ipsi/contra=0.37/0.42; SCR: *d* ipsi/contra = 0.41/0.42; PCR *d* ipsi/contra=0.46/0.45, p_FDR_<0.05), cingulum adjacent to the hippocampus (CGH: *d* ipsi/contra=0.40/0.36, p_FDR_<0.05), and the superior fronto-occipital fasciculus (SFO: *d* ipsi/contra=0.31/0.43, p_FDR_<0.05). The fornix was the only structure that exhibited a higher MD effect size within the younger TLE cohort (*d*=0.41, p_FDR_<0.05) than in the older TLE cohort (*d*=0.27, p_FDR_<0.05). Apart from the fornix, all other regions exhibited significantly increased effect for the older TLE cohort compared to their age-matched controls. Group-by-age interaction analyses for both FA and MD of the white matter structures showed almost all regions were significantly more increased for MD and decreased for FA in patients than their healthy control counterparts (p_FDR_<0.05).

The trajectories of FA measures across age (**Fig. 2**) resembled grey matter trajectories, with a gradual downward trend until age 50, followed by a steeper decline. This general trend occurs across most white matter tracts, such as the limbic, cortical-subcortical, and cortical structures (see Table **S2** for a complete list). The splenium, body, and genu of the corpus callosum white matter tracts were more pronounced in their FA decreases. Likewise, MD age trajectories for all white matter tracts gradually increased until age 60, after which MD values showed even greater increases. Most pronounced findings were with structures associated with the limbic, cortical-subcortical, and cortical regions bilaterally. The correlation of the age trajectories with the TLE-MTS subcohort again were again much higher (FA/MD *r*=0.86/0.78) compared to their TLE-NL sub-cohort counterparts (FA/MD *r*=0.57/0.50). Additional correlations calculated between the FA/MD age trajectories showed a much more positive correlation with the right-sided TLE patient cohort (FA/MD *r*=0.91/0.86) as opposed to the left-sided patient cohort (FA/MD *r*=0.68/0.75).

**Figure 2.**
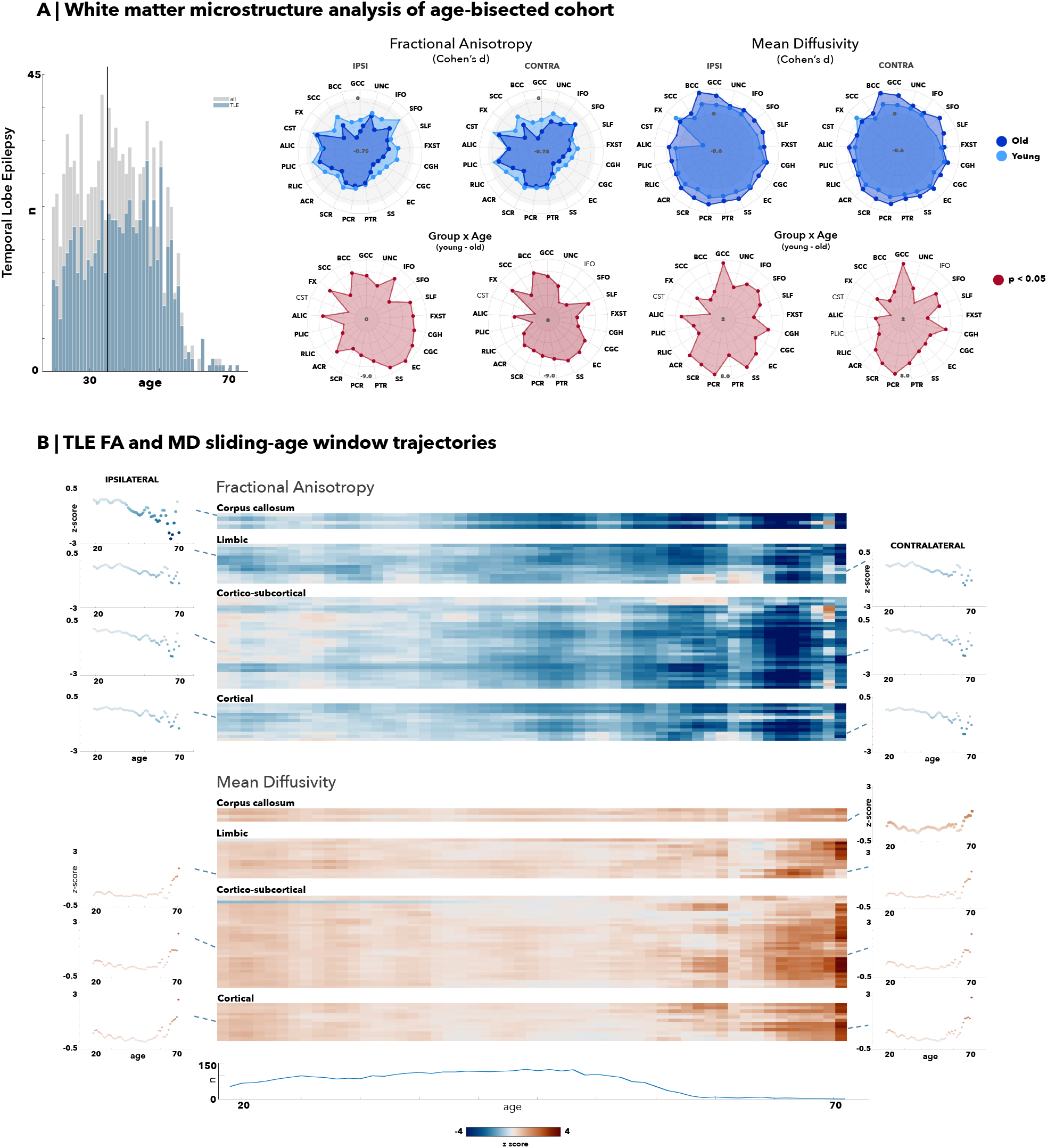
White matter alterations via median-split analysis and sliding-age window trajectories in temporal lobe epilepsy (TLE). **(A)** Median age of 35 was chosen as the division into young and old cohorts. All right-sided temporal lobe epilepsy (TLE) patients were flipped, with results demonstrating almost all white matter tracts significantly decreased for fractional anisotropy (FA) measures and increased for mean diffusivity (MD) measures for both young and old TLE cohorts against their age-matched controls. **(B)** Average age trajectories were calculated for ipsilateral and contralateral white matter tracts associated with the corpus callosum, limbic, cortical-subcortical, and cortical regions. All tracts demonstrate an accelerated and pronounced decline in FA measures past age 50 and a similarly pronounced increase in MD measures past age 60.

### Age-mediated structural covariance dynamics

Aside from region-specific correlations to itself, there is a general positive correlation between FA, SV, and CT measures, and a general negative correlation between MD, SV, and CT measures (**Fig. 3**). This is further illustrated by the scatter plots of every patient and their *z-*scores against controls for grey and white matter measures. The top 10% of correlations were calculated for FA by CT, FA by SV, and CT by SV correlograms, while the bottom 10% of correlations were calculated for MD by CT, MD by SV, and MD by FA to reflect their general trend. In the FA by CT correlogram, more than half of FA structures associated with the right caudal anterior cingulate neocortical region belong in the top 10%. Likewise, more than half of all CT structures correlated with the fornix cres/stria terminalis (FXST), ipsilateral and contralateral posterior thalamic radiation (PTR) were found in the top 10%. The ipsilateral posterior thalamic (PTR) radiation was again implicated in the FA by SV correlogram. Finally, in the SV by CT correlogram, the right pallidum had almost all its correlations with the neocortical regions within the top 10% of connections. Similar analyses done within the MD by CT, MD by SV, and MD by FA implicate the MD value of contralateral corticospinal tract (CST) for both FA white matter regions and subcortical volumes. Both ipsilateral and contralateral FXST were also strongly associated across most neocortical and subcortical regions. Covariance patterns between the ipsilateral/ipsilateral, ipsilateral/contralateral, and contralateral/contralateral structures showed concordant trends across hemispheres (**Fig. S4)**.

**Figure 3.**
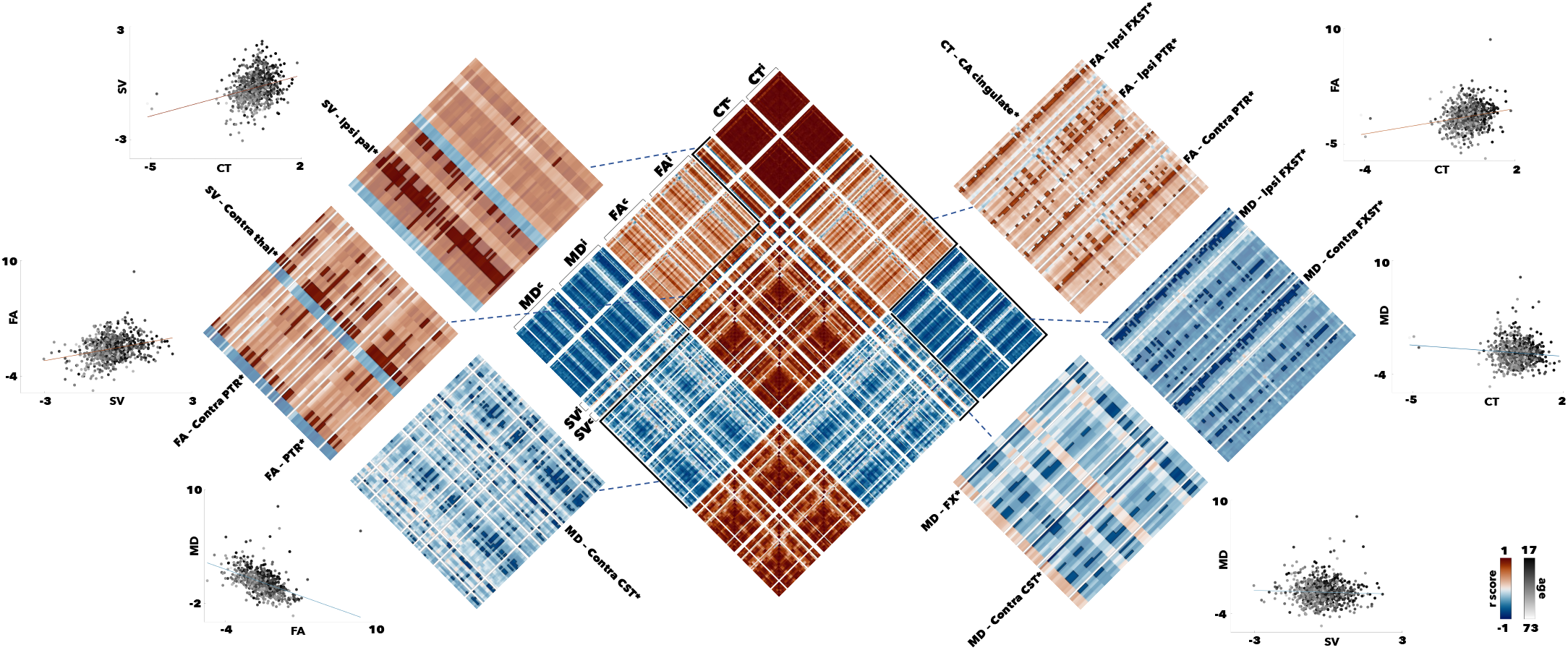
Sliding age-window structural covariance correlations in temporal lobe epilepsy (TLE). Correlograms (center) showing the correlational of each region-associated age trajectory across the lifespan for CT, SVC, FA, and MD were calculated. The top 10% (*red*) and bottom 10% (*blue*) of connections were preserved to identify regions more closely associated with each grey or white matter measures. Scatter plots demonstrate the age-dependent (*grey*) distribution of z-score values against controls.

## DISCUSSION

Our multi-site cross-sectional study investigated aging effects on brain structural compromise across the lifespan in pharmaco-resistant TLE. Benefiting from a large sample size, multiparametric assessment of grey and white matter alterations, and standardized protocols for data extraction and processing from all ENIGMA-Epilepsy sites, our findings illustrate TLE as a temporo-limbic network disorder exhibiting *(i)* age-mediated decline of grey/white matter structural integrity, *(ii)* an acceleration of effects with advanced age (>55 years). Collectively, these cross-sectional findings motivate future longitudinal studies that comprehensively track epilepsy related effects across the lifespan and underscore the need for prompt diagnosis and treatment in patients suffering from pharmaco-resistant seizures.

In our cohort, the MRI-derived grey matter atrophy became more pronounced in older patients and extended over a broader anatomical territory with advancing age. Prior literature on cortical thickness, hippocampal volumes, and subcortical structures supports a pattern of diffuse whole brain atrophy in TLE,^6,9,14,16,18^ echoing our findings of reductions in the entorhinal, medial parietal, cingulate, and frontal cortices as well as hippocampal and thalamic volumes. Other cross-sectional cohort studies from ENIGMA-Epilepsy have also highlighted broad cortical atrophy and subcortical volume reductions in TLE patients on a group level.^6,9^ However, little has been done to investigate the influence of age on such alterations. Previous evaluations of cortical thinning across the lifespan in healthy individuals have revealed that the entorhinal and temporopolar cortices seem to increase initially following an inverse U-shape pattern, unlike most other neocortical regions where cortical thickness is highest during childhood.^29^ Thus, our findings of linearly progressive thinning, particularly for the entorhinal and temporopolar regions, is in line with cumulative disease effects and complement previous reports of disease progression in epilepsy.^16^ Furthermore, previous studies often only take age into account when assessing disease progression in TLE patients. They tend to report no correlations between age and grey matter atrophy despite compelling evidence for decline in normal aging in adulthood,^14,15,19,30^ or identify age as a likely determinant on atrophy and volume loss.^17^ A meta-analysis reviewing progressive atrophy in TLE has shown that most prior reports are cross-sectional with narrow age-ranges, small cohort sizes, and consequently, poor age control.^16^ Our age-stratified atrophy analysis demonstrates that age has increasing influence through later years in patients compared to controls, and may thus need to be incorporated in biomarker development and validation efforts. The increased rate of cortical thinning and subcortical volume reductions of TLE patients older than 50 years is similar to previous longitudinal studies demonstrating increased progression and severity of cortical thinning in TLE cohorts,^14,15^ reflecting an increased vulnerability to cortical structural compromise in later decades of life.

In line with prior investigations showing white matter alterations in TLE,^8,31–33^ diffusion MRI derived metrics across almost all white matter tracts were altered in our young and old TLE cohorts compared to age-matched controls at the group level. MD increases and FA reductions were most marked bilaterally in the fornix, cingulum, and corpus callosum. These findings add to prior evidence for a pronounced impact of TLE on temporo-limbic fibre systems, which is notable given their close anatomical and functional links to the mesio-temporal TLE epicentre.^34^ Accelerated structural changes in temporo-limbic tracts across the lifespan is compatible with secondary damage, consistent with previous studies on adult TLE that showed distinct MD alterations in early-onset TLE patients.^35^ Furthermore, the microstructural perturbations of almost all white matter fascicles suggests whole-scale white matter compromise of anatomical networks, particularly in older TLE patients. Nevertheless, the relationship between white matter atrophy and aging remains inadequately explored in TLE; most earlier studies on group level differences or spatial characteristics indicate no correlation between age and white matter metrics due to limited cohort sizes,^33,36^ entirely omit age analyses,^37^ or report significant associations of disease duration after correcting for age.^8,31^ Age correction alone does not, however, eliminate all age-mediated effects on disease duration or age of onset. Older patients, even if diagnosed more recently, may still exhibit an accelerated rate of gray and white matter loss compared to their younger counterparts. Given the correlation of age, age of onset, and disease duration, future work should precisely account for the confounding between these variables.

Our work also includes a novel evaluation of inter-regional covariance patterns of white and grey matter changes over the lifespan. In TLE, covariance approaches have previously been restricted to single compartments, primarily the cortical grey matter.^10,38^ Here, we extended this approach by simultaneous analyses of grey and white matter compartments, and by cross-correlating age-related change curves. Our results implicate the fornix, posterior thalamic radiation, corticospinal tracts, pallidum, and thalamus as structures most strongly correlated to the global age-related change curves of at least one other metric (CT, SV, FA, or MD). These findings of overall coupled grey and white matter progression reinforce the notion of TLE as a network disorder, which is likely impacted by pathological processes that affect multiple regions and compartments simultaneously.^36,39^ Limbic-associated cortical regions are known to exhibit greater atrophy in TLE patients, and similar pathology within temporo-limbic white matter fascicles have been previously shown.^33,39^ Considering their internal anatomy, microstructure, and development, temporo-limbic systems have been shown to harbor an elevated potential for brain plasticity and susceptibility to insults in epilepsy.^7^ Notably, temporo-limbic structures are also known to be involved in functional networks implicated in affective and cognitive processes, notably memory.^40^ Thus, our findings suggesting age-dependent alterations in limbic pathways may also help explain the increased cognitive decline as well as affective difficulties often experienced by older individuals with TLE.^13^ More information regarding cognitive, electro-clinical, socio-demographic, and behavioral data in these participants is needed to further ascertain these relationships in future work, which ideally also adopts a harmonized multisite perspective as the current study.

Continued seizures are the reflection of a chronic disease process, and have been associated with progressive grey and white matter compromise in patients,^41,42^ and neuronal damage in experimental models of TLE.^43^ Recent findings also suggest that vascular risk factors, altered tissue perfusion, and neurodegenerative deposits of amyloid and hyperphosphorylated tau in epilepsy may contribute to structural compromise.^44–47^ Beyond the role of these biological and clinical factors, multiple lines of evidence suggest an increasing risk of psychosocial deprivation in older epilepsy patients and of a psychiatric diagnosis.^48,49^ Continued antiseizure medical treatment could also modulate the age-related structural compromise seen here.^50^ These factors likely coalesce to explain the accelerated structural compromise we see in TLE patients, and particularly the older subgroup, but the cross-sectional nature of this present study in evaluating these aging effects lends susceptibility to confounders such as cohort effects and prevents the identification causal associations. A significant proportion of our cohort had paediatric age of onset, therefore more comprehensive lifespan analyses could begin around the mean age of onset to fully illustrate grey and white matter trajectories. Stratified sampling and increased recruitment of older age populations given our limited cohort size past age 55 would also reinforce the argument for accelerated disease processes linked to aging as opposed to cohort effects. Finally, our analyses did not correct for disease duration as doing so in a cross-sectional study may fail to show discernable differences.^16^

Our findings indicate that age, whether acting as a surrogate for disease duration or used to argue for disease progression, is a clinically highly relevant factor in TLE, particularly in patients above the age of 55. The heterogeneity of older patients in educational attainment, socioeconomic status, or other factors that confer protective effects on cortical thinning and cognitive outcomes may necessitate an individualized approach and more comprehensive data to evaluate the true risk baseline age poses for accelerated brain atrophy and potential worsening of symptoms. In this context, future longitudinal work with stratified sampling across the lifespan is recommended to explore epilepsy-specific characteristics such as frequency of seizures alongside additional contributions of neurological, neurovascular, and psychiatric comorbidities. Longitudinal designs may also help to between address cohort-level health effects such as differences in drug exposure, shifts in treatment, and world-scale events. Ultimately, future work should be directed towards patients-specific longitudinal tracking of network-level trajectories to fully comprehend the extent of disease progression, its potential acceleration in older age, and associations to functionally and clinically relevant outcomes.

## Supporting information

Supplemental_all

## ACKNOWLEDGEMENTS

R.R.C. is funded by the Fonds de la Recherche du Québec – Santé (FRQS). L.C. is supported by the Berkeley Fellowship jointly awarded by UCL and Gonville and Caius College, Cambridge, and by Brain Research UK (award 14181). A.B. and N.B. are supported by FRQS and CIHR. B.C.B. acknowledges research support from the NSERC Discovery-1304413, CIHR (FDN-154298, PJT-174995), SickKids Foundation (NI17-039), the Helmholtz International BigBrain Analytics and Learning Laboratory (HIBALL), HBHL, Brain Canada, and the Tier-2 Canada Research Chairs program. G.P.W. acknowledges support from the MRC (G0802012, MR/M00841X/1). C.R.M. acknowledges research support from the NIH/NINDS (R01 NS124585; R01 NS122827). P.S. acknowledges support from the PNRR-MUR-M4C2 PE0000006 Research Program “MNESYS”. IRCCS ‘G. Gaslini’ is a member of ERN-Epicare. S.L. acknowledges funding from the Canadian Institutes of Health Research (CIHR). E.B. has been partially supported by grant-RC and the 5x1000 voluntary contributions, Italian Ministry of Health. F.C., C.L.Y, and M.K.M.Y. acknowledges support from FAPESP (Sao Paulo Research Foundation) grant number 2013/07559-3.

