## Supplemental_all for "A worldwide ENIGMA study on epilepsy-related gray and white matter compromise across the adult lifespan"

### SUPPLEMENTARY

#### Table S1 | Detailed clinico-demographic data of ENIGMA-Epilepsy TLE cohort

| **Case-control cohort** | **Age** (years) | **Sex** (F/M) | **Age of onset** (years) | **Disease duration** (years) | **Engel score** (I) |
|---|---|---|---|---|---|
| HC ($n$ = 886) | 36.0 ± 11.7 | 508/378 | – | – | – |
| L TLE MTS ($n$=280) | 39.0 ± 10.8 | 161/119 | 14.9 ± 11.8* | 23.8 ± 13.5* | 102 |
| R TLE MTS ($n$=226) | 39.8 ± 10.9 | 130/96 | 15.1 ± 11.3* | 24.4 ± 13.8* | 66 |
| L TLE non-lesional ($n$= 150) | 36.0 ± 10.5 | 93/133 | 19.4 ± 11.4* | 16.8 ± 12.0* | 23 |
| R TLE non-lesional ($n$=113) | 365.5 ± 11.5 | 71/42 | 19.9 ± 13.4* | 14.8 ± 10.9* | 25 |

Age, age of onset, and disease duration are presented in mean ± SD. *Clinical information available: a) age of onset: L-TLE MTS = 8/280, R-TLE MTS = 11/130, L-TLE non-lesional = 8/150, R-TLE non-lesional 7/113; b) disease duration: L-TLE MTS = 32/280, R-TLE MTS = 24/130, L-TLE non-lesional 16/150, R-TLE non-lesional 18/113.

**TABLE S2 | White matter tract labels**

| Group | Tract abbreviation | Full name |
|---|---|---|
| Corpus Callosum | SCC | Splenium of the corpus callosum |
| | BCC | Body of the corpus callosum |
| | GCC | Genu of the corpus callosum |
| Limbic | FX | Fornix (column and body) |
| | FXST | Fornix (cres) / Stria terminalis |
| | CGH | Cingulum (hippocampus) |
| | CCG | Cingulum (cingulate gyrus) |
| | UNC | Uncinate fasciculus |
| Cortical-subcortical | ALIC | Anterior limb of the internal capsule |
| | PLIC | Posterior limb of the internal capsule |
| | RLIC | Retrolenticular part of the internal capsule |
| | ACR | Anterior corona radiata |
| | SCR | Superior corona radiata |
| | PCR | Posterior corona radiata |
| | PTR | Posterior thalamic radiation |
| | EC | External capsule |
| | CST | Corticospinal tract |
| Cortical | SS | Sagittal stratum |
| | SLF | Superior longitudinal fasciculus |
| | SFO | Superior fronto-occipital fasciculus |
| | IFO | Inferior fronto-occipital fasciculus |

**FIGURE S3 | Sensitivity analyses with age-sliding window ranges**

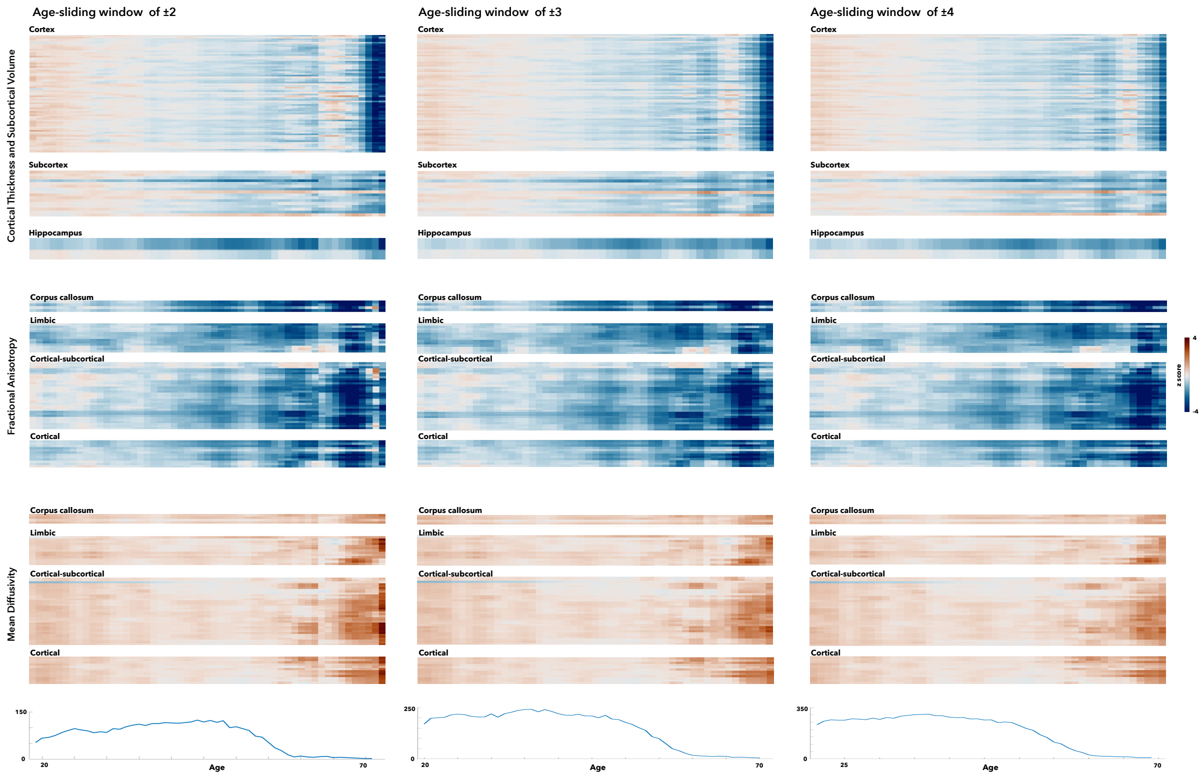

**FIGURE S4 | Covariance of lifespan trajectories separated by ipsilateral and contralateral structures and regions**

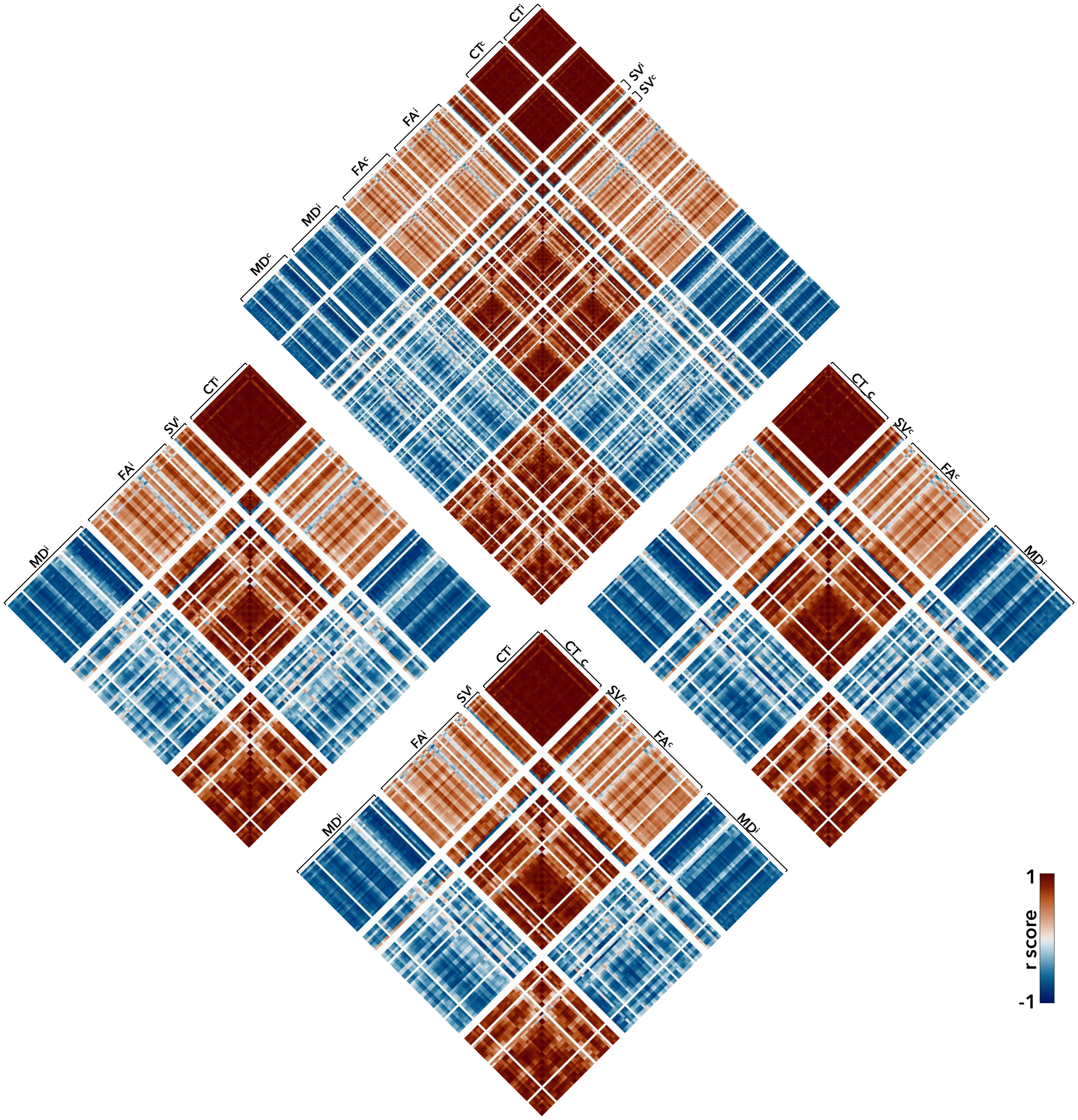
